# P2X7-dependent exchange of extracellular microparticles and mitochondria by mouse microglia

**DOI:** 10.1101/2023.04.18.537325

**Authors:** Simonetta Falzoni, Paola Chiozzi, Valentina Vultaggio-Poma, Mario Tarantini, Elena Adinolfi, Paola Boldrini, Anna Lisa Giuliani, Dariusz C. Gorecki, Francesco Di Virgilio

**Author notes:** Correspondence to Dr Francesco Di Virgilio, Department of Medical Sciences, University of Ferrara, Via Borsari 46, 44121 Ferrara, Italy.

## Abstract

Microparticles (MPs) are ubiquitously secreted by all cells and play a fundamental role in numerous biological processes such as cell-to-cell communication, cell differentiation, inflammation, and cell energy transfer. Ligation of the P2X7 receptor (P2X7R) by extracellular ATP (eATP) is the well-known stimulus for MP release, which affects their contents in a cell-specific fashion. We investigated MP release and functional impact in mouse microglial cell lines characterized for high (N13-P2X7R^High^) or low (N13-P2X7R^Low^) expression of the P2X7R. Stimulation with extracellular ATP triggered a P2X7R-dependent release of a MP population enriched with naked mitochondria. Released mitochondria were taken up and incorporated into the mitochondrial network of the recipient cells in a P2X7R-dependent fashion. Other constituents of the MP cargo, e.g. NLRP3 and the P2X7R itself, were also delivered to the recipient cells. Transfer of mitochondria, NLRP3 and P2X7R increased the energy level of the recipient cells and conferred a pro-inflammatory phenotype. These data show that P2X7R-dependent exchange of MPs and mitochondria modulates energy metabolism and inflammatory responses, pointing to the P2X7R as a master regulator of intercellular organelle and MP trafficking in mouse microglia.

## Introduction

Information exchange is a vital function of all living organisms, whether uni- or multi-cellular. Neurotransmitters, hormones, growth factors, and inflammatory mediators are usually released and travel across the intercellular space as individual molecules, but often they are packed in the lumen of exosomes/vesicles/particles, collectively referred to as microparticles (MPs), to be delivered to nearby or faraway target cells ^1^. While exosomes originate from cytoplasmic multivesicular bodies, extracellular vesicles are the result of plasma membrane budding, thus they usually contain cytoplasmic components, including mitochondria ^2, 3^. Mechanism of MP release has been the focus of intense scrutiny over the last two decades, leading to the identification of the crucial role played by extracellular ATP (eATP) acting at the P2X7 receptor (P2X7R) ^4–8^. It is an established fact that eATP is an ubiquitous messenger accumulating at sites of trauma, cancer or inflammation, and also acting as a modulator of neurotransmission in the central nervous system (CNS) ^9 10^. Effects of eATP are mediated by the well-known P2 receptor family, notably at inflammatory and cancer sites by the P2X7R subtype ^11^. Verderio and co-workers showed previously that astrocyte-derived eATP causes vesicle shedding from microglia ^5^. We showed a similar effect of eATP in human dendritic cells ^6, 7^. The potent MP-releasing activity of eATP acting at the P2X7R has become textbook knowledge ever since ^12^. This eATP effect is of special interest because, besides promoting MP release, this nucleotide also modulates MP contents, and is itself carried by the MPs ^8, 13^. The discovery that MPs contain functioning mitochondria that they can transfer to the target cells, has added an additional level of interest and complexity to the pathophysiological function of these extracellular structures ^2^.

Naked mitochondria have been identified in the mixed vesicle population collectively referred to as shed MPs ^8, 14^. We recently observed that eATP stimulation of mouse melanoma tumor cells via the P2X7R triggers a massive release of mitochondria-laden MPs as well as of naked functional mitochondria ^8^. The function of extracellular free mitochondria is still debated, nonetheless there is a good evidence that these organelles undergo cell-to-cell transfer via tubular structures ^15^, MP uptake ^16^, or phagocytosis, and can be incorporated into the target cell mitochondrial network. Mitochondria transfer is increasingly recognized as a key process in tissue repair ^17^, in CNS homeostasis ^18^, and in immune-cancer cell interaction, as a determinant of immune evasion ^19^. While macrophages appear to be a leading participant in the process of mitochondrial transfer, both as donors and recipients ^20^, much less is known about that role in microglia. In the CNS, there is good evidence that mitochondria transfer occurs between astrocytes and microglia and bi-directionally between neurons and astrocytes ^21^, but only scattered observations report an inter-microglia mitochondria transfer ^22^. Exchange of mitochondria between neurons and glia, or between microglial cells, is thought to be relevant in a multiplicity of neuroinflammatory and neurodegenerative diseases, and thus potentially exploitable for therapeutic purposes.

In the present study we set to investigate how P2X7R expression affects mitochondria release and microglia-to-microglia mitochondria transfer, and whether it modulates the MP cargo. To address this issue, we took advantage of the N13 microglia cell line expressing high levels of the P2X7R (N13-P2X7R^High^) and of a variant selected in our laboratory for low P2X7R expression (N13-P2X7R^Low^). We found that P2X7R expression on the donor cells strongly affects both basal and eATP-stimulated MP release, as well as MP uptake by the target cells. The MP fraction released upon eATP stimulation contained a heterogenous population of small vesicles, mitochondria-containing vesicles and naked mitochondria. Upon interaction with the target cell, mitochondria and MP components (including the P2X7R itself) were transferred to the recipient cells. MPs derived from N13-P2X7R^High^ were very efficiently transferring the P2X7R to N13-P2X7R^Low^, thus enabling the restoration of P2X7R-dependent responses including reversible plasma membrane permeabilization, the hallmark of P2X7 function. Furthermore, the ability to generate multinucleated giant cells (MGCs), typical histological features of chronic inflammation, was also restored. MP-mediated transfer between microglial cells might be an efficient mechanism for the modulation of neuroinflammation.

## Results

Resting or stimulated microglial cells are known to release a mixed population of MPs of different sizes containing both exosomes and plasma membrane-derived extracellular vesicles ^5, 12^. Accruing evidence supports a key role of the P2X7R in MP release in microglia as well as in other cell types^6, 7, 23, 24^. N13-P2X7R^High^ and N13-P2X7R^Low^ microglial cells were loaded with the cytoplasmic marker dyes calcein/AM (Fig. 1a-d) or carboxyfluorescein succinimidyl ester (CSFE) (Fig. 1e-h), then rinsed and incubated in the absence or presence of either eATP or the more potent analogue benzoyl-ATP (Bz-ATP). As shown in Fig. 1, N13-P2X7R^High^ cells (Fig. 1a, b), but not N13-P2X7R^Low^ (Fig. 1c, d), released MPs into the culture medium within few minutes of incubation in the presence of the P2X7R agonists. Challenge with eATP (Fig. 1e) or Bz-ATP (Fig. 1f), up to a concentration of 1 mM or 0.5 mM, respectively, significantly increased MP release from N13-P2X7R^High^ cells, several fold and time-dependently (Fig. 1h), while MP release from N13-P2X7R^Low^ was negligible. MP release was linearly related to the eATP but not to the Bz-ATP concentration, likely because BzATP concentrations higher than 0.3 mM cause necrotic cell death. Both eATP- and Bz-ATP-stimulated MP release was mitigated, albeit not fully inhibited, by pretreatment with the potent covalent P2X7R blocker oxidized-ATP (oxo-ATP) (Fig. 1g), as previously shown ^25^.

**Fig. 1.**
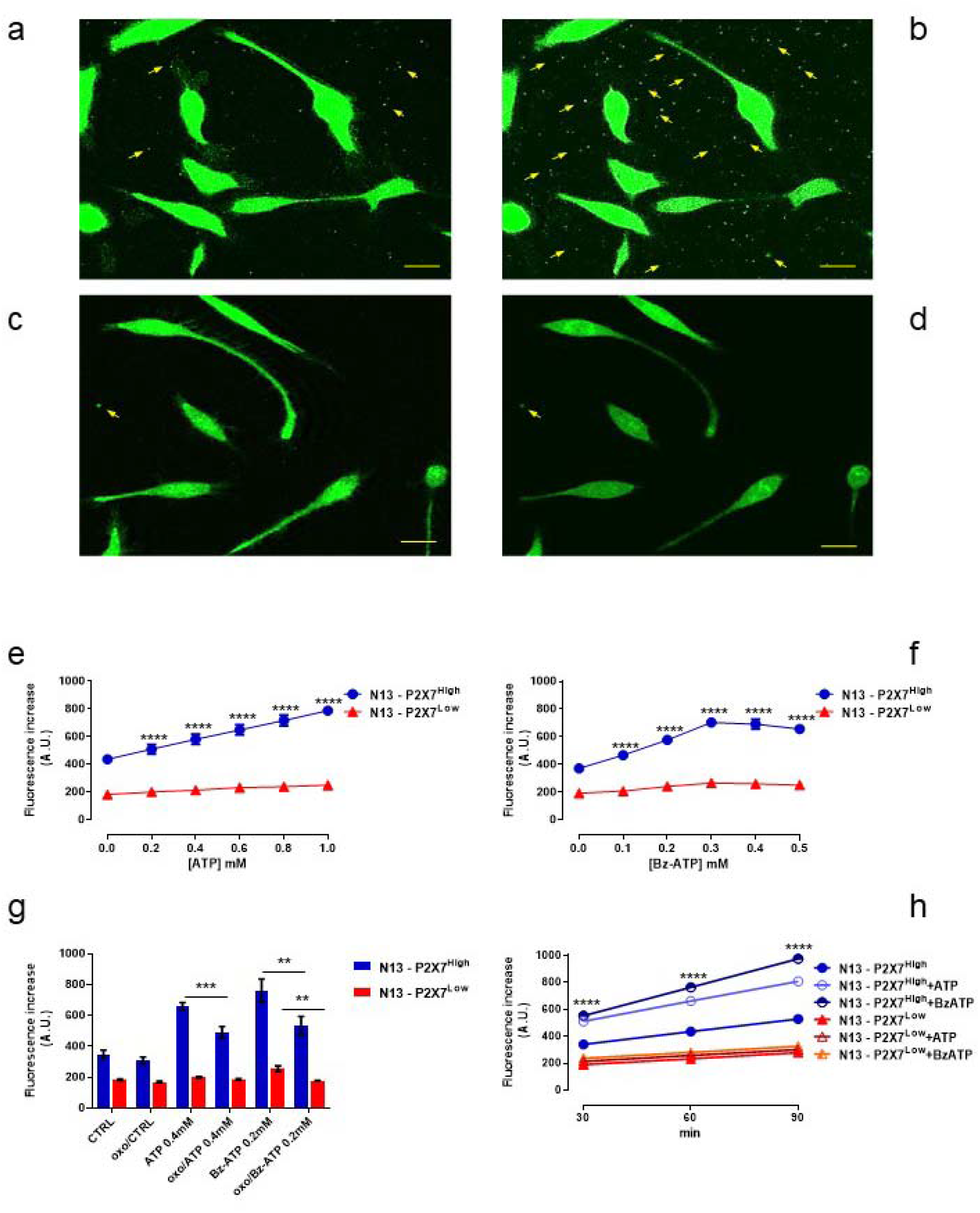
N13 microglial cells release MPs in a P2X7R-dependent fashion. Five x10^5^ N13-P2X7R^High^ (**a)** or N13-P2X7R^Low^ (**c)** cells were loaded with the cytoplasmic marker calcein/AM (1 μg/mL) in RPMI complete culture medium for 20 min, then rinsed, re-suspended in the same medium and challenged with 0.4 mM eATP for 15 min to trigger MP (arrow) release (**b**, eATP-stimulated N13-P2X7R^High^; **d**, eATP-stimulated N13-P2X7R^Low^). **e-h**, cells were loaded with CSFE (5 μM), and incubated with increasing concentrations of eATP (**e**) or BzATP (**f**) for 60 min. **g**, cells were treated with 0.3 mM oxidized-ATP (oxo) for 1 h at 37° prior to stimulation with either eATP or BzATP. **h**, Time course of MP release in the presence or absence of either 0.4 mM eATP or 0.2 mM BzATP. To measure MP fluorescence, cell supernatants were withdrawn, centrifuged at 10,000x*g* (4 °C) for 1 h to concentrate the MPs, and the pellet resuspended in 500 μL of sucrose medium. Fluorescence was measured in a fluorimeter cuvette at the 488/520 nm wavelength pair. Data are means ± SEM of three to four experiments each performed in triplicate. ** = p<0.01; *** = p<0.001; **** = p<0.0001 by Student -test (**e**-**g**), or two ways ANOVA (**h**). Fluorescence images were acquired with a Nikon Eclipse TE 300 inverted fluorescence microscope equipped with a 40X objective and fluorescein filters. Bar = 10 μm.

N13-P2X7R^High^ cells were then labelled with the selective mitochondrial marker Mitotracker Green FM, and left unchallenged (Fig. 2a) or stimulated with eATP (Fig. 2b). Besides intracellular organelles, also extracellular MPs were stained by this dye, suggesting that they contain mitochondria (Fig. 2a, b, arrows). To further characterize the mitochondrial content, MPs were isolated, as described in Materials and Methods, from the supernatants of resting (Fig. 2c, d), or eATP-stimulated (Fig. 2e, f) N13-P2X7R^High^ microglia, incubated in the presence of either calcein/AM (Fig. 2c, e) or the selective, potential-sensitive, mitochondrial stain Mitotraker Red FM (Fig. 2d, f), and analyzed for fluorescence emission. Henceforth, MPs released from N13-P2X7R^High^ or N13-P2X7R^Low^ microglia will be referred to as MP-P2X7R^High^ or MP-P2X7R^Low^, respectively. As shown in Fig. 2g, calcein-positive MPs are about twice as many as those positive for Mitotracker Red, suggesting that either no all calcein-positive MPs contain mitochondria, or that calcein- and MitoTracker Red-positive extracellular vesicles are separate MP populations. To verify this latter hypothesis, we co-labeled the source N13-P2X7R^High^ cells with both calcein/AM and MitoTracker Red FM prior to eATP challenge. Image merging showed that quite a few MPs were yellow, indicating co-localization and suggesting that the MP fraction contained a mixed population of naked mitochondria, MPs containing mitochondria and mitochondria-less MPs (Fig. 2h).

**Fig. 2.**
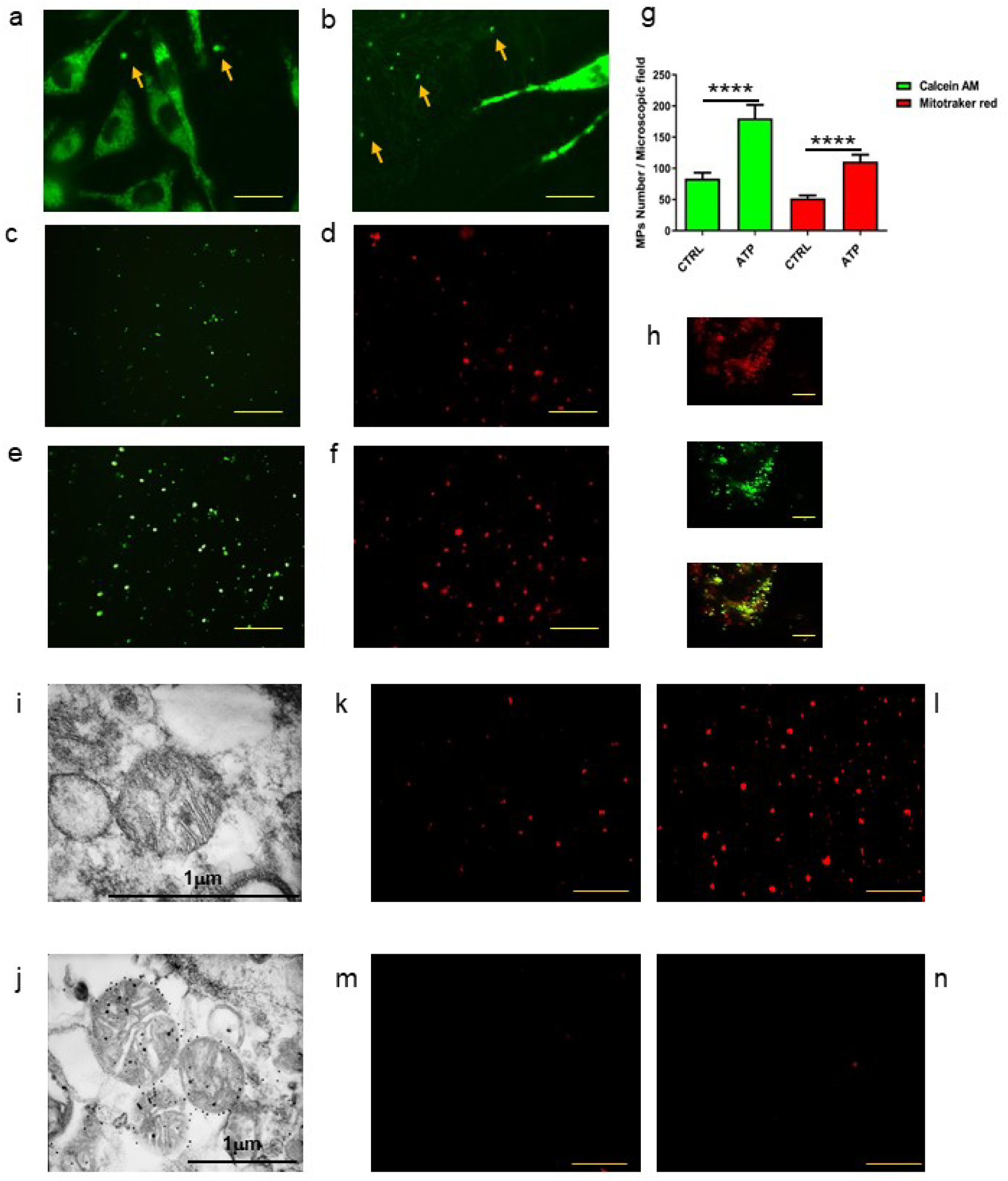
Released MPs contain mitochondria. N13-P2X7R^High^ (**a, b**) cells (10^5^) were seeded onto 13 mm diameter round glass coverslip and stained with the mitochondrial selective dye MitoTracker Green FM (200 nM) for 15 min at 37°C. The coverslips were then placed on the heated (37° C) stage of a Nikon inverted microscope as described in the legend to Fig. 1, and either left unchallenged (**a**) or stimulated with 0.4 mM eATP for 5 min (**b**). Yellow arrows indicate MitoTracker Green FM-stained MPs. In panels **c-f**, supernatants from resting (**c, d**) or 0.4 mM eATP-stimulated (**e, f**) N13-P2X7^High^ were stained with calcein/AM (**c, e**) or MitoTracker Red FM (**d, f**) and analyzed with a Zeiss LSM510 confocal microscope equipped with a 63X oil immersion plan-apochromat objective. **g**, calcein/AM- or MitoTracker Red FM-positive MPs released from resting or eATP-stimulated N13-P2X7^High^ released over a 60 min timespan were quantitated. **h**, MPs released over a 24 h incubation period from eATP (0.4 mM)-challenged N13-P2X7R^High^ cells co-labeled MitoTracker Red FM (200 nM) and calcein/AM (1 μM), were isolated as described in Materials and Methods and analyzed by confocal microscopy with rodhamine (upper) or fluorescein (middle) filters, and merged (lower). MPs isolated from N13-P2X7R^High^ were analyzed by TEM without (**i**) or with (**j**) immunogold labeling with an antibody against the specific mitochondrial protein TOM20. MPs isolated from resting (**k**) or 0.4 mM eATP-stimulated (**l-n**) N13-P2X7R^High^ were stained with TMRM and analyzed by confocal microscopy. 2.5 μM FCCP (**m**), or 200 nM rotenone (**n**) were also added. Bars = 20 μm (**a-f**), 10 μm (**h, k-n**). Data are means ± SEM from three independent experiments each performed in triplicate. **** p < 0.0001 by unpaired t test.

To further verify the presence of mitochondria, MPs were examined by transmission electron microscopy (TEM) and labelled with immunogold using an antibody against a specific transporter of the outer mitochondrial membrane (TOM20). MP-P2X7R^High^, recovered from the supernatant of eATP-stimulated N13-P2X7R^High^ cells, contained naked mitochondria (Fig. 2i) that were specifically labelled by the anti-TOM20 antibody (Fig. 2j). Thus, as previously shown by us and by other investigators ^2, 8, 26^, MPs are a rather heterogenous population containing a plethora of intracellular molecules and even intracellular organelles, including mitochondria. MPs, whether released spontaneously (Fig. 2k) or upon eATP stimulation (Fig. 2l-n), were also strongly stained by the potential-sensitive dye tetramethylrhodamine methyl ester (TMRM), a further proof of a membrane potential, negative inside. Accordingly, addition of the mitochondrial uncoupler carbonyl cyanide-p-trifluoromethoxyphenylhydrazone (FCCP) (Fig. 2m), or of the electron transport chain inhibitor rotenone (Fig. 2n) fully abrogated TMRM fluorescence.

As additional characterization, we probed by Western blotting the MP fraction for canonical mitochondrial markers, for the inflammasome protein NOD-LRR- and pyrin domain-containing protein 3 (NLRP3) and for the P2X7R itself. MP-P2X7R^Low^ compared to MP-P2X7R^High^ had a lower content of Cyt C, VDAC1, TOM20, TIM23, as well as of NLRP3 and P2X7R (Fig. 3a-h). Challenge of the N13-P2X7R^High^ or N13-P2X7R^Low^ cells with eATP or BzATP caused a marked increase in the content of mitochondrial markers, P2X7R and NLRP3 in MP-P2X7R^High^, but a significantly lower one in MP-P2X7R^Low^.

**Figure 3.**
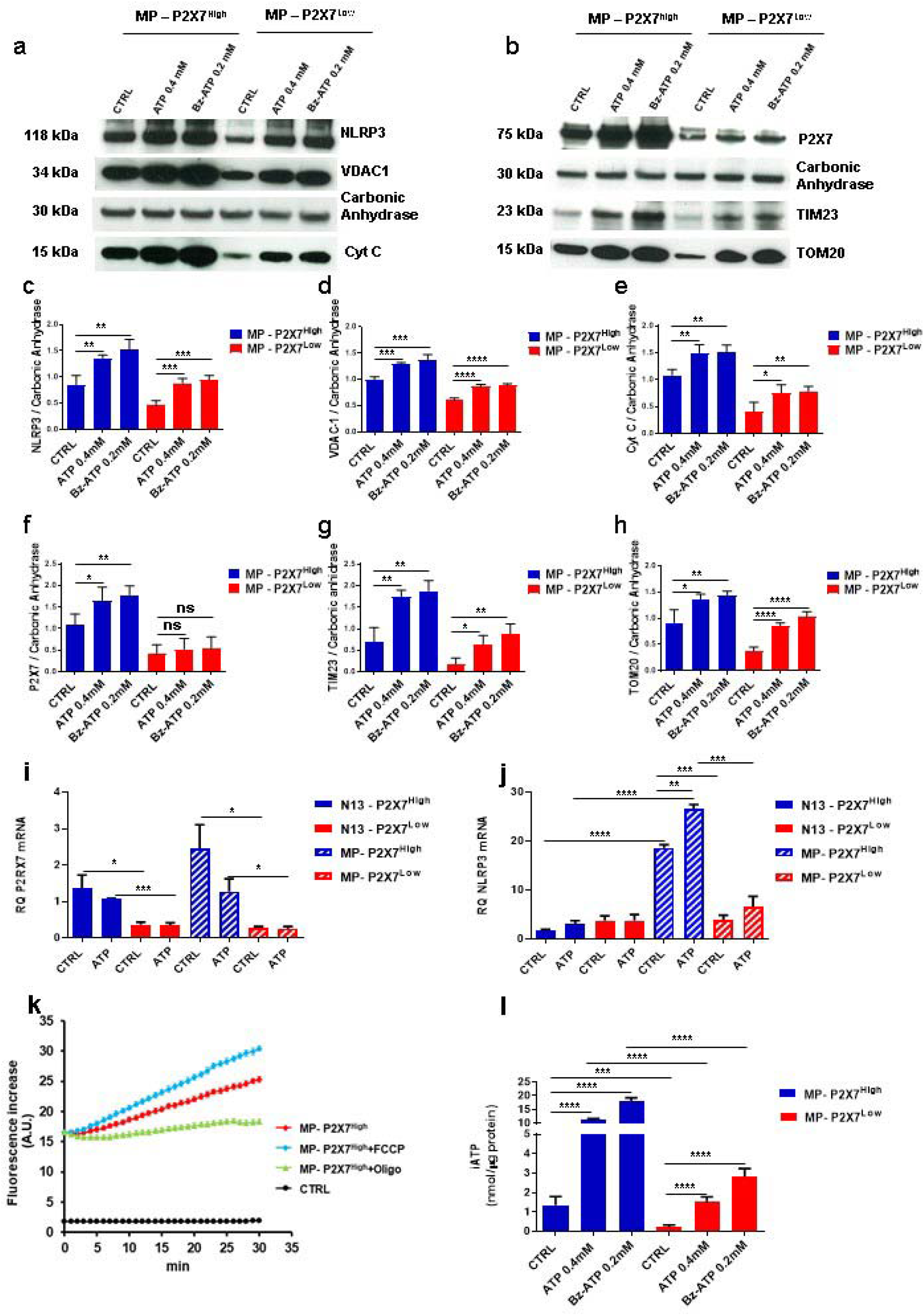
MP-P2X7R^High^ compared to MP-P2X7R^Low^ express higher levels of NLRP3, P2X7R and mitochondrial markers. N13-P2X7R^High^ and N13-P2X7R^Low^ cells were plated in 75 cm^2^ culture flasks and stimulated with 0.4 mM eATP or 0.2 mM BzATP for 24 hours. At the end of this incubation, MPs were isolated and processed for WB analysis (**a, b**) as described in Materials and Methods. Carbonic anhydrase was used as a loading control. Densitometry (**c-h**) was performed with ImageJ software. mRNA was isolated from control or eATP-stimulated (0.4 mM) N13-P2X7R^High^ (**i**) or N13-P2X7R^Low^ (**j**), and from released MPs during a 24 h incubation. CTRL, control resting cells. **k**, Oxygen consumption by MP-P2X7R^High^ in the absence or presence of FCCP or oligomycin. Control (CTRL): O_2_ consumption in the absence of MPs. **l**, intracellular ATP (iATP) content of MP-P2X7R^High^ and MP-P2X7R^Low^. ATP was measured as described in Materials and Methods. Data are means ± SEM from three to four independent experiments. * p < 0.05; ** p < 0.01; *** p < 0.001; **** p < 0.0001 by unpaired t test.

Besides NLRP3 and P2X7R proteins, MPs also contained their respective mRNAs (Fig. 3i, j). Challenge of N13-P2X7R^High^ with eATP had a differential effect on the NLRP3 and P2X7R mRNA content in both of the source cells and in the released MP-P2X7R^High^, with the former being increased and the latter decreased (Fig. 3i, j). Again, changes in NLRP3 or P2X7R mRNA levels in the eATP-stimulated MP-P2X7R^Low^ were negligible.

Despite the identification of mitochondria in the MP fraction by morphological, biochemical and immunolabelling techniques, and demonstration of a FCCP-sensitive membrane potential, the formal proof that mitochondria in the MP fraction were able to support respiration and associated oxidative phosphorylation (OxPhos) was still required. Thus, we investigated whether MPs support O_2_ consumption sensitive to canonical mitochondrial poisons such as FCCP and oligomycin. We were unable to use the SeaHorse technology since it was impossible to make MP adhere to the bottom of SeaHorse dishes without damaging the MPs themselves, thus we turned to a fluorometric technique based on O_2_ quenching of the Abcam reagent (see Materials and Methods). MPs exhibited a basal oxygen consumption that was further stimulated by FCCP and fully blocked by oligomycin, thus confirming its dependency on ATP synthesis (Fig. 3k). In agreement with the higher mitochondrial content, unstimulated MP-P2X7R^High^ accumulated an about 10-fold higher intracellular ATP (iATP) content compared to MP-P2X7R^Low^ (Fig. 3l). The iATP content of MP-P2X7R^High^, and to a lesser extent that of MP-P2X7R^Low^, was further enhanced by stimulation with eATP or Bz-ATP of the respective source cells prior to the MP collection (Fig. 3l).

We next investigated the fate of the released MPs, to verify whether, as reported in other cell types, they might be taken up by target microglial cells. Donor N13-P2X7R^High^ or N13-P2X7R^Low^ cells (5x10^5^) were labelled with Mitotracker Green FM. Then, MPs released over a 24 h incubation were recovered as described in Materials and Methods, and the same dose of MPs administered to unlabeled N13-P2X7R^High^ or to unlabeled N13-P2X7R^Low^ “target” cells. Unlabeled N13-P2X7R^High^ cells were incubated for 24 h with Mitotracker Green FM-labeled MP-P2X7R^High^, rinsed and then analyzed (Fig. 4a). This prolonged co-incubation caused a strong staining of intracellular vesicular and filamentous structures in unlabeled N13-P2X7R^High^. Alternatively, unlabeled N13-P2X7R^Low^ cells were incubated with Mitotracker Green-labeled MP-P2X7R^High^. In this case, staining of intracellular structures was much weaker. Then, unlabeled N13-P2X7R^High^ cells were challenged with Mitotracker Green-labeled MP-P2X7R^Low^, and finally N13-P2X7R^Low^ were challenged with Mitotracker Green-labeled MP-P2X7R^Low^. In both these latter cases, staining of intracellular organelles was very weak, albeit slightly stronger when N13-P2X7R^High^ were challenged with MP-P2X7R^Low^.

**Fig. 4.**
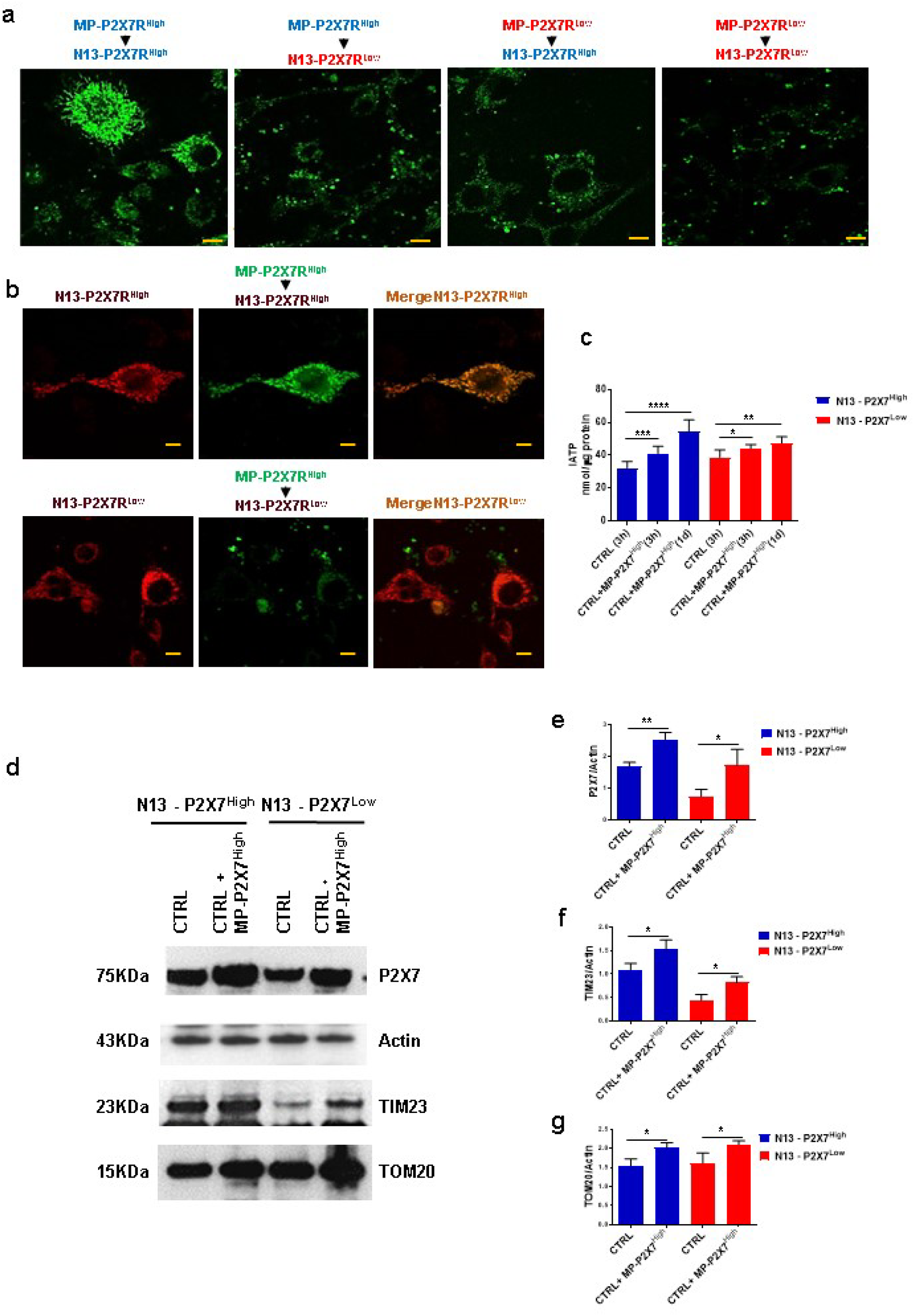
Released mitochondria are taken up by target microglial cells in a P2X7R-dependent fashion. **a**, unlabeled N13-P2X7R^High^ or N13-P2X7R^Low^ were let adhere to 24-well dishes, rinsed and further incubated in complete RPMI medium for 24 h with MitoTracker Green FM-labeled MP-P2X7R^High^ or MP-P2X7R^Low^ previously isolated from MitoTracker Green FM-labeled source N13 cells (see Materials and Methods). At the end of this incubation, the monolayers were rinsed to remove extracellular MPs, fresh medium was added and the cells analyzed by confocal microscopy as described in the legend to Fig. 1. **b**, N13-P2X7R^High^ (upper panel) or N13-P2X7R^Low^ (lower panel) were loaded with MitoTracker Red FM (400 nM) at 37 C° for 20 min, then thoroughly rinsed and further incubated at 37° C in complete RPMI medium for 24 h with MP-P2X7R^High^ released from MitoTracker Green-stained N13-P2X7R^High^. At the end of this incubation period cells were analyzed by confocal microscopy. **c**, intracellular ATP (iATP) content of N13-P2X7R^High^ or N13-P2X7R^Low^ after 3 h or 1 d of co-incubation with MP-P2X7R^High^. **d**, WB of control or MP-P2X7R^High^-pulsed N13-P2X7R^High^ or N13-P2X7R^Low^ cell lysates. Actin was used as loading control. **e-g**, densitometric analysis. Isolation of MP-P2X7R^High^ and co-incubation with either N13-P2X7R^High^ or N13-P2X7R^Low^ were performed as described in Materials and Methods. **a,** Bar = 20 μm; **b**, bar = 10 μm. * p < 0.05; ** p < 0.01; *** p < 0.001; **** p < 0.0001 by unpaired t test.

To verify that mitochondria in the MP fraction transferred to the target cells were able to fuse with the endogenous mitochondrial network, recipient N13-P2X7R^High^ (Fig. 4b, upper panels) or N13-P2X7R^Low^ cells (Fig. 4b, lower panels) were labeled with Mitotracker Red FM, and co-incubated in the presence of Mitotracker Green FM-labeled MP-P2X7R^High^. Within 6 h, nearly all of the added MPs were taken up by N13-P2X7R^High^ but not by N13-P2X7R^Low^. Image merging showed an almost perfect colocalization with the endogenous, red-stained, mitochondrial network in N13-P2X7R^High^ but far less in N13-P2X7R^Low^ cells (merge). Mitochondria uptake also enhanced the total ATP content of the recipient cells, but especially of the N13-P2X7R^High^, since iATP levels in these cells after a 1 d incubation in the presence of MPs almost doubled (Fig. 4 c). Furthermore, co-incubation of N13-P2X7R^High^ or N13-P2X7R^Low^ with MP-P2X7R^High^ increased the TIM23, TOM20 and P2X7R content of the recipient cells (Fig. 4d-g).

To rule out the possibility that spillover of green or red fluorescence might at least partially confuse the mitochondria uptake and colocalization analysis, we run control experiments with single fluorochrome-labeled cells showing that there was absolutely no spillover of red fluorescence into the green channel and vice versa (Supplementary Fig. S1)

Next, we verified whether the MPs challenge could modify cell responses. Cells with high P2X7R expression exhibit the peculiar reversible permeabilization of the plasma membrane triggered by exposure to eATP ^27–29^, a feature witnessed by lucifer yellow uptake by N13-P2X7R^High^ cells (Fig. 5a). In contrast, N13-P2X7R^Low^ microglia was refractory to eATP-mediated permeabilization, as shown by near absent dye uptake (Fig. 5c). Incubation in the presence of MP-P2X7R^High^ only slightly enhanced the already high lucifer yellow uptake in N13-P2X7R^High^ cells (Fig. 5b), but had a significant impact on this dye uptake in N13-P2X7R^Low^ cells (Fig. 5d). Measurement of cell-associated fluorescence shows that incubation in the presence of MP-P2X7R^High^ almost doubled lucifer yellow uptake by N13-P2X7R^Low^ cells (Fig. 5e). In addition, incubation of N13-P2X7R^High^ cells in the presence of MP-P2X7R^High^ almost doubled their proliferation rate, both at the 48 h and 72 h time points (Fig. 5f). The effect of the incubation of N13-P2X7R^High^ with MP-P2X7R^Low^ was much smaller, albeit still statistically significant (Fig. 5f). N13-P2X7R^Low^ cells showed a lower basal proliferation rate compared to N13-P2X7R^High^, a well-known feature of P2X7R-less cells ^30–32^, and their proliferation was enhanced by incubation with MP-P2X7R^High^ and to a lesser extent with MP-P2X7R^Low^ (Fig. 5g).

**Fig. 5.**
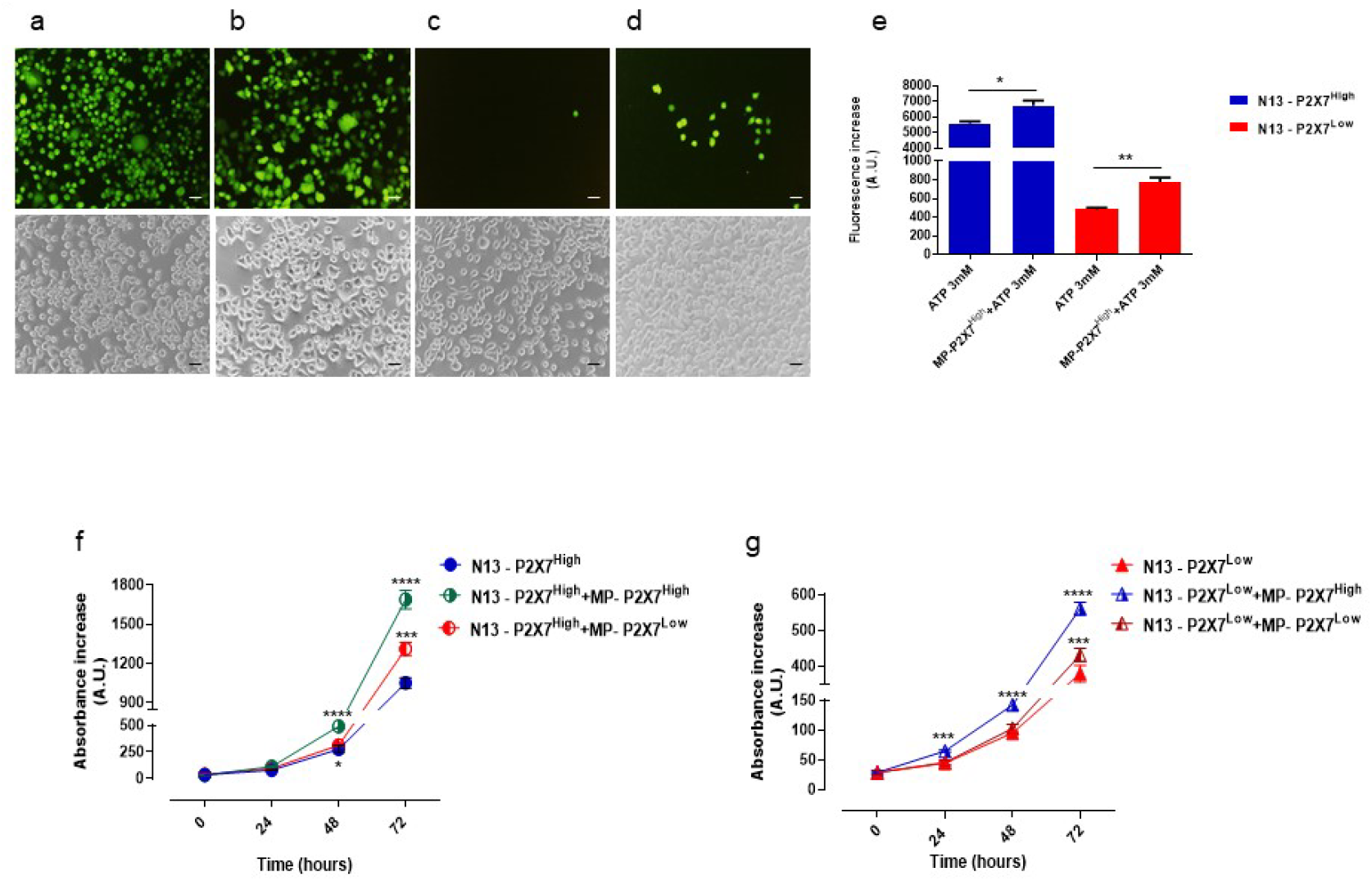
Fusion of MP-P2X7R^High^ with the recipient cells restores eATP-mediated plasma membrane permeabilization and increases the proliferation rate in a P2X7R-dependent fashion. Fluorescence (**a-d**, upper panels) or phase contrast (**a-d**, lower panels) images of eATP-stimulated lucifer yellow uptake of control (**a**) or MP-P2X7R^High^-challenged (**b**) N13-P2X7R^High^, or control (**c**) or MP-P2X7R^High^-challenged (**d**) N13-P2X7R^Low^. Cells were incubated in complete RPMI medium at 37 °C in a CO_2_ incubator in the absence or presence of MP-P2X7R^High^ for 24 h, then rinsed, further incubated in complete RPMI at 37° C and challenged with 3 mM eATP in the presence of 1 mg/mL lucifer yellow. At the end of this incubation, the monolayers were thoroughly rinsed, incubated in fresh complete RPMI medium and analyzed by fluorescence microscopy with a IMT-2 Olympus phase/fluorescence microscope equipped with 20x objective. Lucifer yellow fluorescence was quantitated as described in Materials and Methods. N13-P2X7R^High^ (**f**) or N13-P2X7R^Low^ (**g**) were seeded at 37° C in a CO_2_ incubator in 24-well plastic dishes in complete RPMI medium. After an overnight incubation, a suspension of either MP-P2X7R^High^ or MP-P2X7R^Low^ was added to each given cell type, and incubation carried out at 37° C in a CO_2_ incubator for further 72 h. Cell number was quantitated by crystal violet (see Materials and Methods). Bars = 50 μm. * p < 0.05; ** p < 0.01; *** p < 0.001; **** p < 0.0001 by unpaired t test.

Expression of the P27XR confers several inflammatory features to mononuclear phagocytes, among which is the ability to generate multinucleated giant cells (MGCs) ^33^, typical for chronic granulomatous inflammation. We previously showed that P2X7R-less phagocytes are unable to undergo spontaneous fusion ^33, 34^, thus we investigated whether fusion of N13 microglia was enhanced after incubation with MPs isolated from either N13-P2X7R^Low^ or N13-P2X7R^High^. MGC formation was measured in N13-P2X7R^High^ cultures incubated for 48 h alone (Fig. 6a), or in the presence of MP-P2X7R^Low^ (Fig. 6b), or MP-P2X7R^High^ (Fig. 6c). As shown quantitatively in Fig. 6d, fusion increased over time, to finally decline after 48 h, as previously documented ^33^, likely due to the accelerated necrosis of the MGCs. Fusion was accelerated significantly by co-incubation with MP-P2X7R^High^ but not with MP-P2X7R^Low^ (Fig. 6d), however co-incubation in the presence of MP-P2X7R^High^ also accelerated MGC necrosis (Fig.6c). Contrary to N13-P2X7R^High^, N13-P2X7R^Low^ cells were unable to generate MGCs spontaneously (Fig. 6e), however their fusion was greatly enhanced by the presence of MP-P2X7R^High^ (Fig. 6g), but not MP-P2X7R^Low^ (Fig. 6f). The strong potentiation of MGC formation promoted by co-incubation in the presence of MP-P2X7R^High^ was quantitated (Fig. 6h).

**Figure 6.**
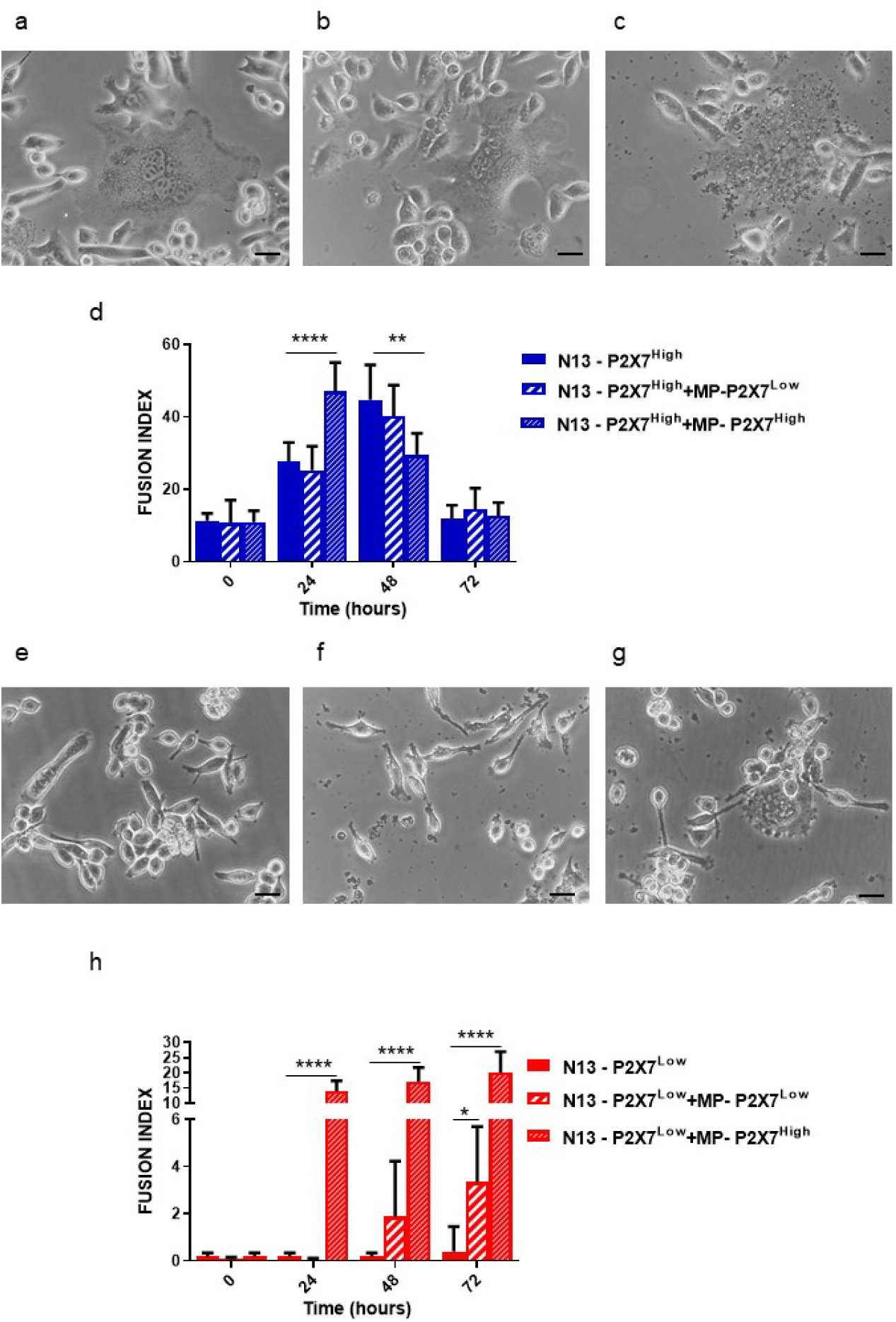
MP fusion with the recipient cells enhances MGC formation in a P2X7R-dependent fashion. N13-P2X7R^High^ were incubated in 24-well plastic dishes at 37° C in complete RPMI medium for 48 h as such (**a**), or in the presence of either MP-P2X7R^Low^ (**b**), or MP-P2X7R^High^ (**c**). Time course of MGC formation shown as fusion index in N13-P2X7R^High^ monolayers incubated in the presence of MP-P2X7R^Low^, MP-P2X7R^High^, or left unchallenged (**d**). N13-P2X7R^Low^ were incubated in 24-well plastic dishes at 37° C in complete RPMI medium for 48 h as such (**e**), or in the presence of either MP-P2X7R^Low^ (**f**), or MP-P2X7R^High^ (**g**). Time course of MGC formation in N13-P2X7R^Low^ monolayers incubated in the presence of MP-P2X7R^Low^, MP-P2X7R^High^, or left unchallenged (**h**). Fusion index was calculated as follows: (number of nuclei within MGC/total number of nuclei) X 100. All pictures were taken with an Olympus microscope as described in the legend to Fig. 5 ** p < 0.01; *** p < 0.001; **** p < 0.0001 by unpaired t test.

## Discussion

We show here that the P2X7R is a master regulator of exchange of functioning mitochondria and MP by microglia. Intercellular transfer of mitochondria via different routes (tunnelling nanotubes, MPs, connexins, uptake of secreted naked mitochondria) is a novel mean of information transfer active in most tissues, including epithelia, neuronal cells, immune and cancer cells ^35^. This mechanism appear to fulfil different needs, such as disposal of damaged mitochondria but also refurbishment of stressed cells with healthy mitochondria. This may have important pathophysiological implications, as the supply of cancer cells with functioning mitochondria has been shown to enhance tumor ability to escape immune surveillance ^19^. Furthermore, transfer of potent damage-associated molecular patterns (DAMPs) such as mitochondrial DNA and cytochrome C is inherently prone to ignite or amplify inflammation.

In the brain, an organ particularly susceptible to mitochondrial dysfunction and thus heavily affected by mitochondriopathies, the mitochondrial transfer might have a very important pathophysiological role. Likewise mitochondria are implicated in neuroinflammation ^36^. Accruing evidence shows that astrocytes transfer mitochondria to neurons ^17^, and vice versa that damaged mitochondria are transferred from neurons to astrocytes ^18^. Microglia is also able to exchange mitochondria with astrocytes ^37^ or with other microglial cells ^22^. The rationale for such exchange activity seems to be on one hand the acceleration of dysfunctional mitochondria disposal and on the other the propagation of pro-inflammatory signals. Of relevance, preliminary results by Watson et al, published in BioRχiv, September 2021, https://doi.org/10.1101/2021.09.01.458565, suggest that microglia is highly effective in the *in vitro* and *in vivo* transfer of mitochondria to glioblastoma tumors, thus increasing tumorigenecity. This is in line with the observations by Sengupta and co-workers showing the potent cancer-promoting effect of mitochondria transfer from immune to cancer cells ^19^.

Macrophages are heavily involved in the transcellular mitochondria exchange both as donor and recipient cells. This has been shown to occur in the adipose tissue, lungs, heart and likely in the bone marrow, and via several mechanisms, i.e. phagocytosis, tunnelling nanotubes or extracellular vesicle uptake ^20^. Although intercellular mitochondria transfer also occurs in the CNS, the mechanism of mitochondria release and uptake by microglia has been little investigated.

Mitochondria transfer is a powerful mean for the modulation of inflammation. Uptake of stem cell-derived mitochondria by macrophages promotes M2 polarization and down modulates inflammation ^38^. Furthermore in the adipose tissue under physiological conditions an intense intercellular mitochondria trafficking occurs between adipocytes and macrophages ^39^. Paradoxically, the main product of mitochondrial metabolic activity, i.e. ATP, is also a powerful and omnipresent inflammatory mediator ^40, 41^ and a most potent stimulant of extracellular vesicle (or MP) release from brain microglia ^5^. This seems to be an integrated pro-inflammatory system as eATP acting at P2X7R, a plasma membrane receptor highly expressed by mononuclear phagocytes, microglia included, promotes MP release ^29, 42^.

The negligible MP release from N13-P2X7R^Low^ reported in the present study confirms that in microglia P2X7R expression is a near absolute requirement for MP release stimulated by eATP, or by its more potent agonist Bz-ATP. The MP population is highly heterogenous including small (about 100-200 nm) and large (100-1000 nm) MPs as well as naked mitochondria. Expression of the P2X7R on the donor cell heavily affects the MP mitochondria content as MP-P2X7R^Low^ showed a much lower level of all specific mitochondrial markers compared to MP-P2X7R^High^, and accordingly have a lower content of ATP. The reason for the reduced mitochondria content in MP-P2X7R^Low^ might be twofold: on the one hand N13-P2X7R^Low^ have a lower mitochondrial metabolism ^43^ and content compared to N13-P2X7R^High^, on the other mitochondria trapping within the budding MPs or mitochondria extrusion might be less efficient in N13-P2X7R^Low^ microglia. It has been proposed that the dispatch via MPs might be a mean to dispose of malfunctioning mitochondria. However our data show that MP-associated mitochondria are mostly functional. In fact, functionality is a key feature of the transferred mitochondria since their uptake and incorporation in the endogenous mitochondrial network improved the energy metabolism in recipient cells. Indeed, a 24 h incubation of N13-P2X7R^High^ microglia in the presence of MP-P2X7R^High^ caused a near doubling of the iATP content in the recipient cells thus proving that supply of healthy mitochondria increases energy levels in these cells.

The MP cargo includes many bio-active molecules besides intracellular organelles, e.g. NLRP3 and P2X7R proteins as well as their respective mRNAs, thus suggesting that the MP uptake may increase the overall pro-inflammatory activity of the recipient cells. This is clearly shown by the conferment of P2X7R-dependent responses to N13-P2X7R^Low^ cells, possibly due to the direct transfer of P2X7R, that is then directly incorporated into the plasma membrane. Another possibility is the translation of the MP-delivered mRNA. Co-incubation with MP-P2X7R^High^ increased P2X7R protein levels in N13-P2X7R^Low^ cells and made them susceptible to eATP-mediated reversible plasma membrane permeabilization, the hallmark of P2X7R activity. Additional functional responses of recipient cells, such as proliferation, were drastically affected by coincubation with MP-P2X7R^High^. The two-fold increase in proliferation might be due to the increased cell energetics as well as to enhanced P2X7R expression. Supply of functioning (“healthy”) mitochondria was shown to potentiate some key macrophage functions such as M2 polarization and phagocytosis^38, 44^, likely to occur due to the increased iATP content. Also increased P2X7R expression due to MP delivery might concur in the stimulation of cell growth, since basal activation of this receptor has trophic/growth-promoting effect ^30^.

Among the most striking albeit often neglected properties of mononuclear phagocytes, microglia included, is their ability to undergo extensive cell fusion to generate MGCs, a hallmark of chronic granulomatous inflammation ^33^. Ability to undergo spontaneous fusion in the absence of stimulation with inflammatory cytokines is fully dependent on P2X7R expression ^33, 45^. Incubation with MP-P2X7R^High^ enhanced fusion in N13-P2X7R^High^ cultures and more interestingly conferred this ability to N13-P2X7R^Low^ that are normally unable to fuse. This is yet another demonstration of the efficient MP-mediated transfer of bioactive molecules, and of their ability to promote a range of inflammatory responses.

To achieve optimal transfer of the MP cargo, expression of P2X7R on the plasma membrane of both the recipient and donor cells is needed. This might be due to the well-documented fusogenic effect of the P2X7R present on the membranes of both interactors ^33^, or possibly due to a higher phagocytic activity of P2X7R-expressing cells. Furthermore, the P2X7R is known to promote phagocytosis and enhance pinocytosis in microglia ^46, 47^, thus increasing uptake of the MP cargo.

In conclusion, these data show that the P2X7R is a key determinant of MP trafficking in mouse microglia being needed both on the donor and recipient cells. P2X7R-expressing MPs efficiently transfer their mitochondrial cargo to the target cells thus enhancing their energetics and modulating their key functionalities. Given the role of eATP and P2X7R in inflammation these findings help to better understand the complex modulation of microglia functions in neuroinflammation and neurodegeneration.

## Material and Methods

### Cell culture

N13-P2X7R^High^ and N13-P2X7R^Low^ cells were grown in RPMI-1640 medium (Sigma-Aldrich, St. Louis, MO, USA) supplemented with 10% heat-inactivated FBS (Gibco, Thermofisher Scientific, Waltham, MA, USA), 100 U/ml penicillin and 100 μg/ml streptomycin (both from Sigma-Aldrich, cat. # P4333) (complete medium). In some experiments, sucrose medium containing 300 mM sucrose, 1 mM K_2_HPO_4_, 1 mM MgSO_4_, 5.5 mM glucose and 20 mM HEPES (pH 7.4) was used.

### Microparticle (MP) preparation

Five x 10^5^ cells were seeded into 6 well plates in complete medium and cultured overnight. Cells were then rinsed and further maintained in fresh complete medium in presence or absence of 0.4 mM ATP for 24h in a CO_2_ incubator at 37° C. Then, the cell supernatant was withdrawn and centrifuged at 800x*g* for 10 min to remove floating cells and cell debris. MPs were then isolated by centrifugation at 10,000x*g* for 1h at 4°C.

### Staining with CellTrace^TM^ CSFE, calcein/AM, MitoTracker Green or MitoTracker Red

Cells (5x10^5^) or MPs thereof were incubated in complete RPMI medium in the presence of: CellTrace^TM^ CSFE (5 μM) (ThermoFisher Scientific, cat. # C34554), calcein/AM (1μg/ml) (Thermo Fisher Scientifc, cat. # C1430), MitoTracker Green FM (200 nM) (Thermo Fisher Scientifc, cat. # M7514) or MitoTracker Red FM (400 nM) (Thermo Fisher Scientific cat. # M22425) in a CO_2_ incubator for 20 minutes at 37°C. At the end of this incubation cells were rinsed and resuspended in fresh complete RPMI medium, while MPs were washed with warm PBS, centrifuged at 10,000x*g* for 30 minutes, and resuspended in 500 μL of either complete RPMI or sucrose medium. Fluorescence was analyzed in a thermostat controlled (37°C) and magnetically-stirred Cary Eclipse fluorescence spectrophotometer (Agilent Technologies, Milano, Italy) or using a confocal microscope (LSM Carl Zeiss, Oberkochen, Germany) equipped with a plan-apochromat X63 oil immersion objective.

### Oxygen consumption

Oxygen consumption was measured according to the recommended protocol “O_2_ kit Abcam” (cat. # AB197243). Briefly, the MP pellet isolated from 5x10^5^ cells was suspended in 100 μL of sucrose solution supplemented with 0.2 mM CaCl_2_, and transferred to a fluorimeter cuvette containing 400 μL of sucrose solution supplemented with 1mM EGTA to a final volume of 500 μL. This MP suspension was then incubated with 40 μL of Abcam reagent. Oxygen consumption was measured at the 340 nm and 642 nm excitation and emission wavelengths, respectively, in the thermostat-controlled fluorescence spectrophotometer (Agilent Technologies) described above for 30 min. FCCP (2.5 μM), or oligomycin (10 μg/ml) were added as a stimulant or an inhibitor, respectively, of mitochondrial respiration.

### Cell Proliferation Assay

Cell proliferation assay was performed using crystal violet technique. Briefly, 10^4^ cells per well were plated in 12-well plate and cultured at 37° C in a CO_2_ incubator in the presence or absence of MPs isolated from either N13-P2X7R^High^ or N13-P2X7R^Low^ cells stimulated with 0.4 mM eATP. At the 0h, 24h, 48h and 72h timepoints cells were fixed with 4% paraformaldehyde, stained with 0.1% crystal violet, extracted with acetic acid 10%, and analyzed by spectrophotometry (OD 595 nm) in a microplate reader (Wallac Victor3 1420, PerkinElmer, Wellesley, MA, USA).

### Measurement of microparticle and cellular ATP content

ATP was measured in MPs or cell lysates with the luciferase/luciferin method (Enliten rluciferase/luciferin cat. # FF2021, Promega, Italy) in a Firezyme luminometer (Biomedica Diagnostics Inc, Windsor, Canada). Data were expressed as relative luminescence units (RLU) and converted into ATP concentration with an appropriate calibration curve.

### Measurement of microparticle potential

Microparticle membrane potential (ΔΨm) was qualitatively measured by monitoring the uptake of the positively charged TMRM (ThermoFisher Scientific, cat. # I34361) at a concentration of 10 nM by confocal microscopy at an emission wavelength of 570 nm, as previously described ^48^.

### Western Blotting

Cells and MPs were lysed in the following lysis buffer: 300 mM sucrose, 1 mM K_2_HPO_4_, 1 mM MgSO_4_, 5.5 mM glucose, 20 mM HEPES, 1 mM benzamidine, 1 mM phenylmethylsulfonyl fluoride (PMSF), 0.2 μg/mL DNase (cat. # D5025) and 0.3 μg/mL RNase (cat. # R5875) (all Sigma Aldrich). Proteins were separated on NuPage Bis-Tris 4-12% precast gel (Life Technologies, Milano, Italy, cat. # NP0335) and transferred to nitrocellulose membrane (GE Healthcare-Life Sciences, Milano, Italy, cat. # 1060003). After incubation for 1 h with TBS-Tween-20 (0.1%) (Sigma Aldrich, cat. # P9416) supplemented with 2.5% non-fat powdered milk and 0.5% BSA to saturate unspecific binding sites, membranes were incubated overnight with primary rabbit antibodies at 4°C. Anti-TOM20 (Sigma Life Sciences, cat. # HPA011562), anti-TIM23 (ThermoFisher Scientific, cat. # PA5-71877), anti-actin (Sigma Aldrich, cat. # A5060), anti-P2X7R (Merck Millipore, Milano, Italy cat. # AB5346), and anti-carbonic anhydrase II (Sigma Aldrich, cat. # SAB2900749) polyclonal antibodies were used at the 1:1000 dilution. Membranes were incubated with a secondary goat anti-rabbit HRP-conjugated antibody (ThermoFisher Scientific, cat. # A16096) at the 1:3000 dilution for 1h at room temperature. Blots were analyzed by enhanced chemiluminescence (ECL) (Thermofisher Scientific, cat # A38555).

### Transmission electron microscopy (TEM) of isolated microparticles

The MP pellet obtained as described was fixed with 1% paraformaldehyde and 1.25 % glutaraldehyde in 0.15 M cacodylate buffer, postfixed in 2% osmium tetroxide, dehydrated by sequential passages in increasing acetone concentrations and included in Araldite Durcupan ACM (Fluka, Sigma-Aldrich, cat. # 44613). Ultrathin sections were prepared with the Reichert Ultracut S ultramicrotome, counterstained with uranyl acetate in saturated solution and lead citrate according to Reynolds ^49^, and analyzed with a Hitachi H800 transmission electron microscope at 100 Kv (Hitachi High Technologies Corporation, Brughiero, Italy).

### Immunogold labelling of isolated microparticles

Microparticles were fixed in 0.4% paraformaldehyde and permeabilized with 0.05% Triton X-100 at room temperature in PBS for 20 minutes followed by incubation at room temperature in 2% BSA supplemented PBS for 20 minutes to block nonspecific binding. MPs were then incubated overnight with the primary anti-TOM20 rabbit polyclonal antibody (Sigma Life Sciences, cat. # HPA011562) diluted 1:100 in 0.2% BSA-containing PBS. Microparticles were thoroughly washed in PBS and bound antibodies were detected with protein A coated with 20-nm gold at the final dilution of 1:100 in 0.2% BSA-containing PBS. Finally, MPs were rinsed in PBS and processed for electron microscopy.

### Lucifer yellow uptake

N13-P2X7R^High^ or N13-P2X7R^Low^ were seeded in 24-well plastic dishes (20x10^4^ per well) in complete RPMI medium for 24 h. Cells were then rinsed and incubated in 300 μL of a saline solution containing 125 mM NaCl, 5 mM KCl, 1 mM MgSO_4_, 1 mM NaH_2_PO_4_, 20 mM HEPES, 5.5 mM glucose, 250 μM sulfinpyrazone, 5 mM NaHCO_3_, and 1 mM CaCl_2_, pH 7.4, supplemented with 1 mg/mL lucifer yellow. ATP was then added at the concentration of 3 mM and the monolayers were incubated in a CO_2_ incubator (37° C). After 15 min, 5 mM MgSO_4_ was added to each well to chelate excess ATP^4-^, the monolayers were rinsed twice with the above saline solution and further incubated in 400 μL of FBS- and phenol red-free RPMI for microscope analysis. To quantitate lucifer yellow fluorescence, cells were lysed with 4 μL of Triton X-100 for 10 min under continuous stirring, supernatants were withdrawn and fluorescence quantitated fluorimetrically at the excitation/emission wavelength pair 428/540. Florescence was normalized on the protein content measured by the Bradford method.

### Real time PCR

Total RNA was extracted from cells or MPs with Trizol Reagent (ThermoFisher Scientific, cat. # 1559626), and purified with the Pure Link RNA kit (Invitrogen, cat. # 12183018A) as per Manufacturer’s instructions. One μg of total mRNA was reverse transcribed with the High Capacity cDNA Reverse Transcription Kit (Applied Biosystem, Carlsbad, CA, USA). Two μL of cDNA were used as a template for qRT PCR. TaqMan probes for expression of P2X7R and NLRP3 genes, and TaqMan probe for G3PDH housekeeping gene were selected from the ready to use gene expression assay (Applied Biosystem). A comparative analysis using the ΔΔCT method was used to quantitate the fold increase of target cDNA relative to N13 cells.

### Statistical analysis

Data were analyzed with the GraphPad Prism 9 software (GraphPad Software, Inc., La Jolla, CA, USA). Statistical significance was calculated with either two-tailed Student’s t or two ways Anova test, assuming equal SD and variance. All data are shown as mean ± standard error of the mean (SEM). Differences were considered significant at P < 0.05. Coding: *P < 0.05; **P < 0.01; ***P < 0.001; ****P < 0.0001.

## Data availability

The data of this study are available as reasonable consultation with the corresponding authors.

## Acknowledgments

The work described in this study was supported by grants from the Italian Association for Cancer Research (n. IG 13025, IG 18581 and IG 22837) to FDV and EA, the Ministry of Education of Italy (PRIN n. 20178YTNWC), the Cure Alzheimer Fund (USA), the Royal Society Exchange Fellowship IES\R3\170196, PUR-THER Transcan-3 Project STC 2021, and institutional funds from the University of Ferrara. This publication is based upon work from PRESTO COST Action CA21130 supported by COST (European Cooperation in Science and Technology).

## Contributions

SF, PC and V V-P performed most of the experiments and data analysis; MT analyzed part of the data and revised the study; EF and ALG help design the study and revised the text; PB was responsible for TEM; DG and FDV designed the study and wrote the paper.

## Footnotes

‘Supplementary information accompanies the manuscript on the Signal Transduction and Targeted Therapy website http://www.nature.com/sigtrans’

