## Supplemental Figures S1 and S2 for "P2X7-dependent exchange of extracellular microparticles and mitochondria by mouse microglia"

**This PDF file includes:**

Figures. S1 to S2

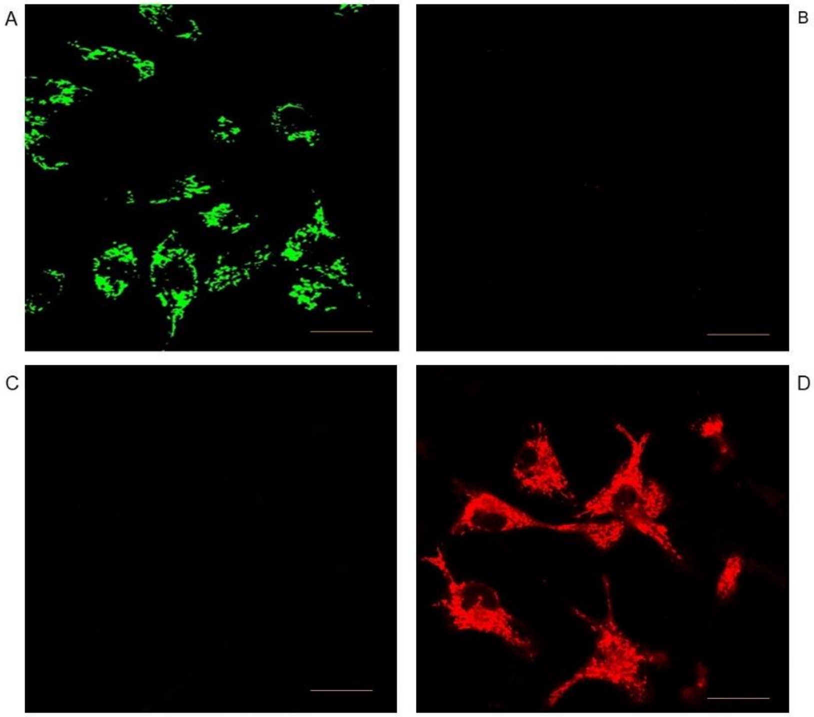


**Fig. S1. Lack of spillover of MitoTracker Green or MitoTracker Red fluorescence.** N13-P2X7R$^{High}$ cells were loaded with MitoTracker Green FM and analyzed with a confocal microscope equipped with either fluorescein (A) or rhodamine filters (C). N13-P2X7R$^{High}$ cells were loaded with MitoTracker Red FM and analyzed with a confocal microscope equipped with either fluorescein (B) or rhodamine filters (D). Loading with the mitochondrial markers was performed a described in Materials and Methods.

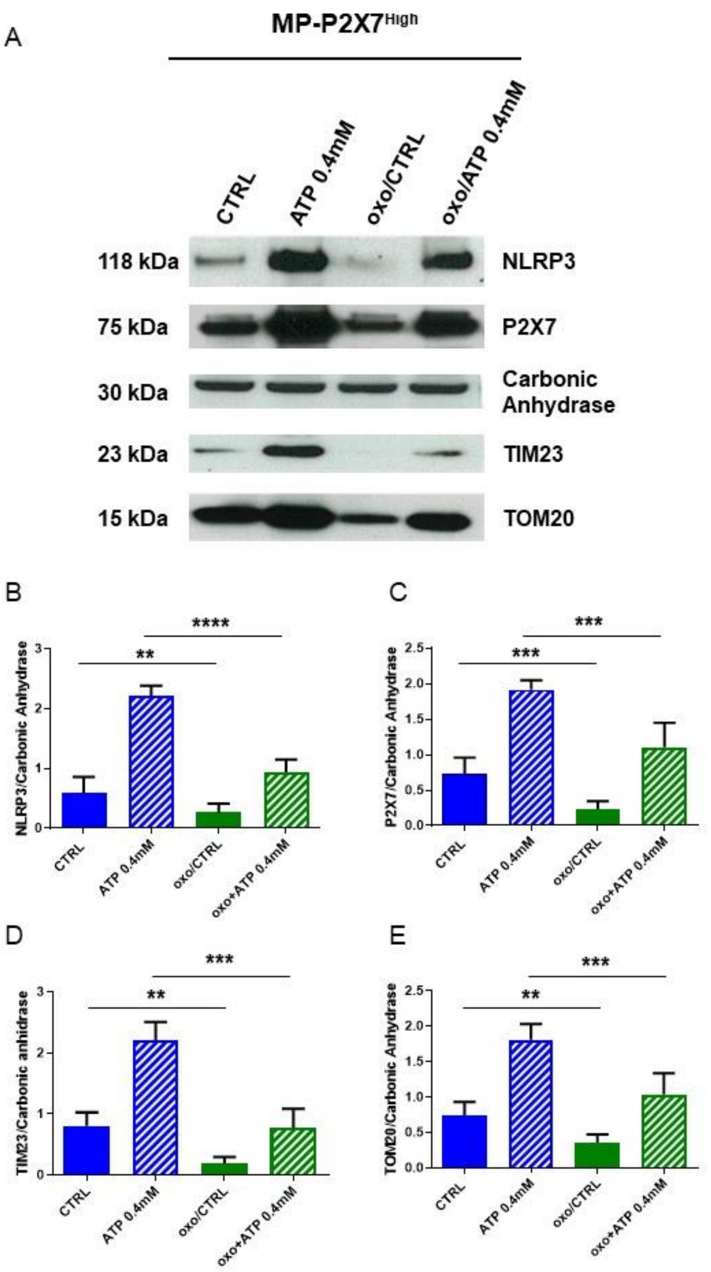


**Fig. S2. The eATP-dependent increase in P2X7R and NLRP3 MP-P2X7R$^{High}$ content is prevented by treatment of the donor cells with oxidized-ATP.** MP-P2X7R$^{High}$ were isolated from N13-P2X7R$^{High}$ cells left unchallenged or treated with eATP (0.4 mM) for 60 min at 37° C in a $CO_2$ incubator. When indicated, cells were pre-treated with 0.3 mM oxidized-ATP (oxo) for 60 min. At the end of the incubaton, cells lysed and analyzed with Western blotting (A) and densitometry (B-E) as described in Materials and Methods. ** $p < 0.01$; *** $p < 0.001$; $p < 0.0001$ by unpaired t test.
